# Hydrophobins from *Aspergillus* mediate fungal interactions with microplastics

**DOI:** 10.1101/2024.11.05.622132

**Authors:** Ross R. Klauer, Rachel Silvestri, Hanna White, Richard D. Hayes, Robert Riley, Anna Lipzen, Kerrie Barry, Igor V. Grigoriev, Jayson Talag, Victoria Bunting, Zachary Stevenson, Kevin V. Solomon, Mark Blenner

**Affiliations:** Chemical and Biomolecular Engineering, University of Delaware, 150 Academy St., Newark, DE 19716; U.S. Department of Energy Joint Genome Institute, Lawrence Berkeley National Laboratory, Berkeley, CA 94720; Department of Plant and Microbial Biology, University of California Berkeley, Berkeley, CA 94720; Arizona Genomics Institute, 1657 E Helen St, Tucson, AZ 85721

**Keywords:** hydrophobin, microplastic, fungi, flocculation

## Abstract

Microplastics present myriad ecological and human health risks including serving as a vector for pathogens in human and animal food chains. However, the specific mechanisms by which pathogenic fungi colonize these microplastics have yet to be explored. In this work, we examine the opportunistic fungal pathogen, *Aspergillus fumigatus*, and other common soil and marine *Aspergilli*, which we found bind microplastics tightly. Up to 3.85+/-1.48 g microplastic plastic/g fungi were bound and flocculated for polypropylene (PP), polyethylene (PE), and polyethylene terephthalate (PET) powders and particles ranging in size from 0.05 – 5 mm. Gene knockouts revealed hydrophobins as a key biomolecule driving microplastic-fungi binding. Moreover, purified hydrophobins were still able to flocculate microplastics independent of the fungus. Our work elucidates a role for hydrophobins in fungal colonization of microplastics and highlights a potential target for mitigating the harm of microplastics through engineered fungal-microplastic interactions.

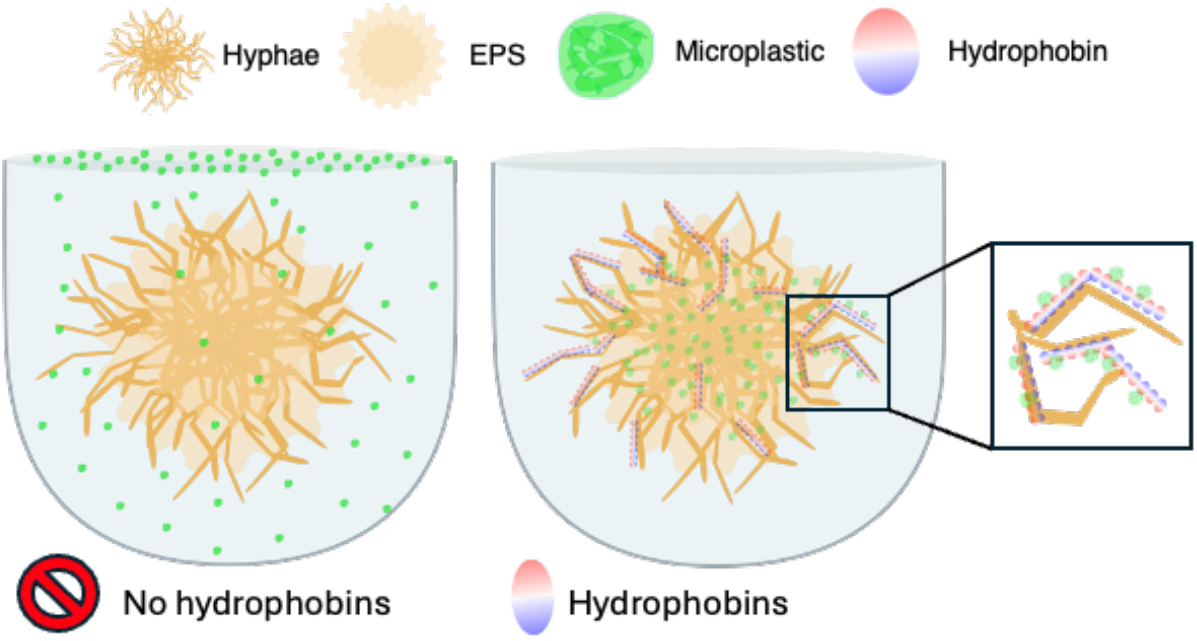

**Significance Statement:** Microplastics pose serious ecological and human health effects by introducing pathogens and toxins into animal and human food chains. Many pathogenic microorganisms preferentially form biofilms on microplastic particles that are then ingested. Here, we demonstrate that hydrophobins, highly hydrophobic, cell surface proteins, enable microplastic binding and colonization by the opportunistic pathogen *Aspergillus fumigatus* and other fungi within the *Aspergillus* genus. Our work recognizes a novel role for hydrophobin proteins, identifying potential strategies for pathogen control and protein-based microplastics recovery.

## Introduction

Microplastics accumulation in the environment has led to myriad ecological and human health issues (1, 2). An estimated 275 million tons of plastic are disposed at end of life each year (3),with the majority of these plastics degraded into microplastic particles dispersed in our air, water and soils (4). As a result of this environmental accumulation, microplastic particles have been observed in the intestinal tracts, tissues, and organs of marine organisms throughout the ecosystem (5). Microplastic particles subsequently work their way through animal and human food chains. Consequently, microplastics have been found in human placentas (6), testis (7), and blood (8) implicating negative health effects from leaching of toxic monomers, additives, and adsorbed environmental pollutants (9, 10). More importantly, polyethylene microplastics in human arteries increased the likelihood of cardiovascular events, stroke, or death, by 2.8-fold relative to a microplastics free control group (11). These environmental and human health issues continue to worsen with exponential increases in plastic production and subsequent increases in environmental contamination (12).

Microorganisms frequently interact with microplastics, forming robust biofilms on their surface. These biofilms are often enriched in pathogens such as those from the genera *Pseudomonas* (13, 14) and *Vibrio* (14). Pathogens tend to be enriched in biofilms due to their ability to promote cell fitness via horizontal gene transfer of antibiotic resistance genes that improve microbial viability of other members of the microbial community (15). Additionally, pathogens such as *Vibrios* have been noted to evolve into hyperbiofilm-formers in stressed microenvironments (15). The presence and enrichment of pathogenic microbes in these biofilms can exacerbate human health impacts by introducing new pathogens into the food chain and harboring increased horizontal transfer of antibiotic resistance genes between pathogens (13, 14, 16). The taxonomic profiles of microplastic-associated biofilms are well documented (17–19), with taxonomic changes to biofilm members vary dependent on sampling location, plastic type, and particle size (17, 20, 21). While there is a strong understanding of the types of microorganisms that bind to microplastic particles under various conditions, the specific biomolecules responsible for microbial binding to microplastics are poorly understood.

Microbes often form biofilms on solid surfaces through secretion of biosurfactants and/or surface proteins. For example, bacteria often rely on flagella or pili to attach to surfaces and form biofilms (22, 23). Additionally, many bacteria secrete extracellular polymeric substances (EPS) containing proteins and lipopolysaccharides (LPS) that promote biofilm hydrophobicity and allow for surface binding (24). Similarly, fungal adhesion to extracellular surfaces is canonically driven by surface proteins called adhesins (25). Adhesins are responsible for cell-cell adhesion, biofilm formation, and adhesion to hydrophobic surfaces in model yeasts like *S. cerevisiae* (26). Common fungi such as *Aspergilli* secrete adhesins belonging to the class hydrophobins that allow them to form strong hyphal networks and adhere to extracellular surfaces (27). Hydrophobins are a class of small (∼10-15kDa), secreted fungal proteins that form amphipathic layers at hydrophobic/hydrophilic interfaces (27), allowing them to bridge fungi to extremely hydrophobic substrates. Though they are known to form strong biofilms on solid surfaces, the interactions of fungi with (micro)plastics are understudied. However, there is growing interest in the fungal members of microplastic-associated biofilms and their interactions due to the inherent pathogenicity of many fungi and their propensity for horizontal gene transfer (20, 28). *Aspergillus niger* has been documented to interact with and bind to polystyrene (PS) and Poly(methyl methacrylate) (PMMA), removing PS and PPMA from solution (29), but binding mechanisms where not studied. These fungi-microplastics relationships are essential to understand how microplastics are colonized, mitigation of health risks from microplastic-bound pathogenic fungi, potential toxicity effects, and how to better remove microbes from microplastics for recovery.

In this study, we leverage *Aspergillus fumigatus*, an opportunistic pathogen found ubiquitously across soil and marine environments (30, 31) to better understand the manner in which fungi bind to microplastics. We isolated a strain of *Aspergillus fumigatus* that forms extremely hydrophobic biofilms, recovering nearly 100% of microplastics from suspensions. Moreover, we confirmed that microplastics recovery occurs ubiquitously across various single and mixed plastic types, confirming that fungal-microplastics interactions are conserved on model post-consumer plastic waste streams. We identified hydrophobin proteins from *A. fumigatus* as the primary driver for microplastic binding by *Aspergillus*. The understanding that hydrophobins are responsible for microplastic binding can be used to reverse biofilm formation by pathogenic strains such as *Aspergillus fumigatus*, subsequently mitigating potential pathogenicity of microplastics. Additionally, hydrophobins can be used in the absence of (pathogenic) hosts to provide sustainable microplastics recovery from aqueous environments.

## Results

### A novel microbial isolate from the yellow mealworm gut microbiome flocculates microplastics from suspension

We discovered a microbial isolate from the gut of *Tenebrio molitor* that flocculates microplastics, pulling them out of suspension. The isolate rapidly (within seconds) flocculated suspended ultra-high molecular weight polyethylene (UHMWPE) particles and floating red fluorescent LDPE particles (Fig. 1A). We evaluated the extent of microplastics flocculation capabilities of this isolate by using 25 mg (0.4% wt./vol) polypropylene (PP), poly(ethylene terephthalate) (PET), surface oxidized UHMWPE, and low-density polyethylene (LDPE) plastics to ensure that the strain can bind to microplastics independent of polymer chemistry and hydrophobicity. The isolate captured 96.0 +/-4.0% of 200 µm LDPE particles, 97.1 +/-0.6% of 50 µm UHMWPE particles, 100 +/-0% of 5 mm PET beads, and 90.9 +/-8.1% of 2 mm PP beads meaning flocculation is independent of both polymer chemistry and particle size (Fig. 1B). Mixed plastic types did not interfere with flocculation. Pairwise combinations of plastics were still recovered with 85-100% efficiency and 92% recovery when a 40 mg (0.8% wt. vol) mixture of all 4 microplastic types and sizes was tested (Fig. 1B). Samples containing PP and PET beads have higher variance due to their larger particle size. If one PP or PET bead was not recovered, it significantly decreases the plastic recovery on the per mass basis. It is likely that the increased surface area of the larger particles requires more binding interactions to retain the plastic within the biofilm, leading to some particles not being captured due to lack of available fungal surface area. Nonetheless, microplastics flocculation is nearly 100% in both ‘pure’ and mixed plastic cases, with pristine and post-consumer plastics of varying chemistries and sizes.

**Figure 1:**
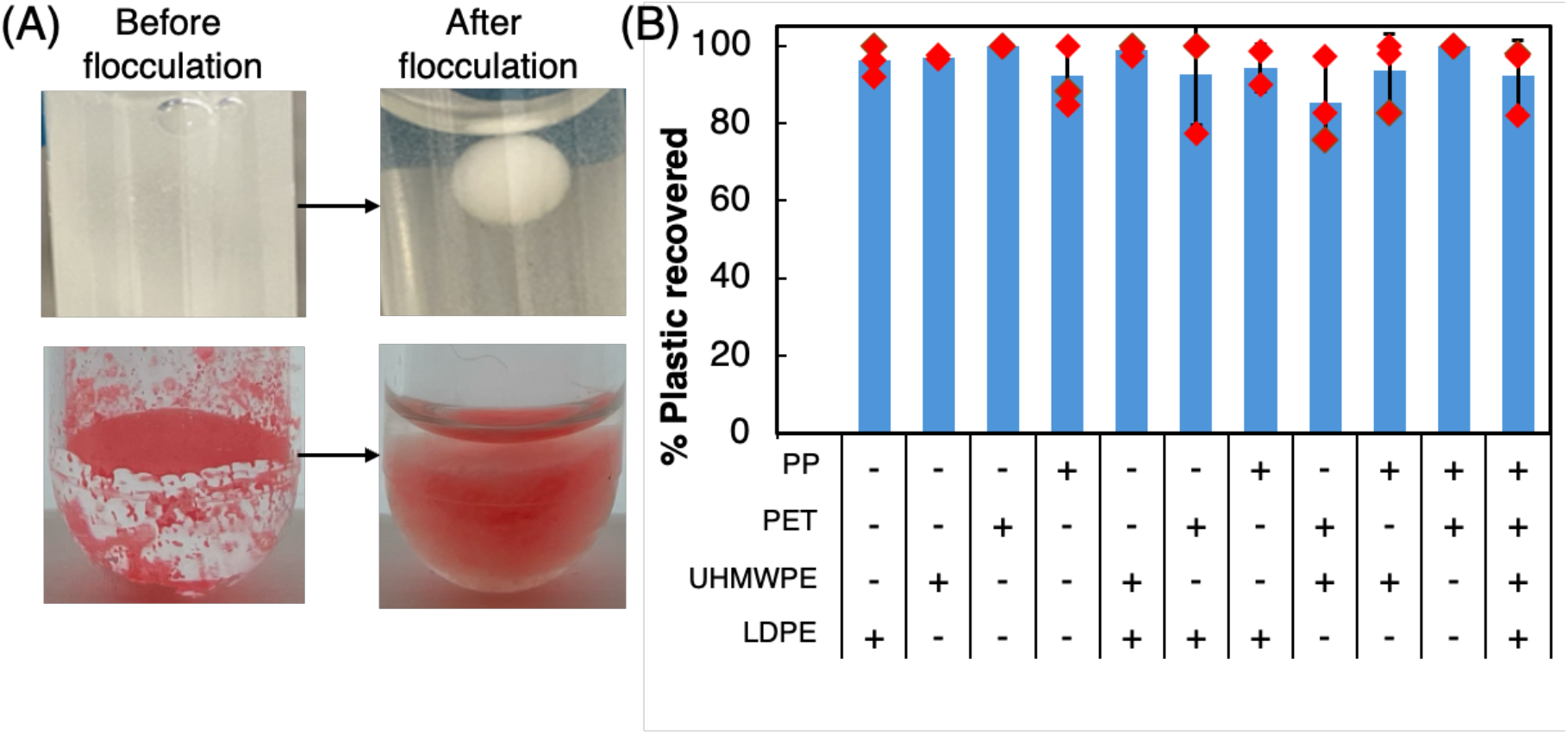
*Aspergillus* ubiquitously capture microplastics from solution. (A) Polyethylene particles captured from solution via flocculation by fungal isolate. (B) Microplastics recovery of a variety of ‘pristine’ and post-consumer plastics shows ubiquitous recovery near 100%.

### Microplastics flocculation is common amongst Aspergillus species

We acquired the whole genome for our microplastic-flocculating isolate and taxonomically placed it as an *Aspergillus* though phylogenetic analysis of its internal transcribed spacer (ITS) (Fig. 2a). The strain was identified as *Aspergillus fumigatus* due to grouping with published *Aspergillus fumigatus* genomes on a species tree constructed using OrthoFinder FastTree (32–36) and was thus named *Aspergillus fumigatus* UD1, hereafter referred to as AF-UD1 (Fig. 2b). Having identified our isolate, we next asked if this ability for microplastics colonization was conserved across the genus by assessing five common *Aspergillus* species spanning a range of phylogenetic distances from AF-UD1 (Fig 2). Each strain successfully flocculated microplastics on an order of 1-5 g of plastic per g of dry biomass (Fig. 3). The recovery of microplastics by all strains implies that there are conserved molecular phenomena occurring in *Aspergilli* cultures that permit microplastics capture.

**Figure 2:**
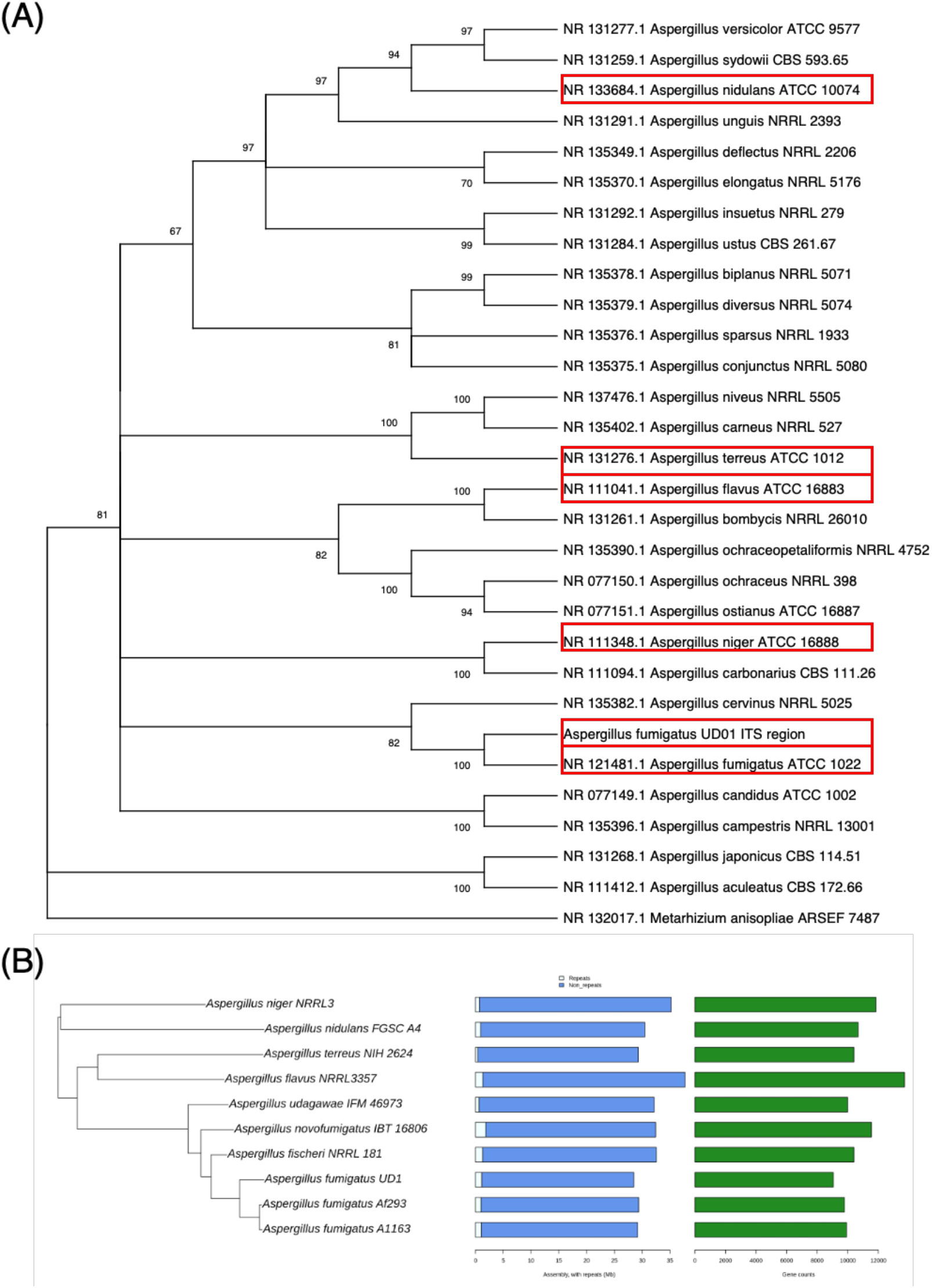
(A) Phylogenetic tree of 45 *Aspergillus* strains built using complete ITS sequences, with strains used in this study boxed in red. A neighbor joining tree was constructed with 100 bootstrap iterations, using *Metarhizium anisopliaei* as the outgroup. Tree is rooted to the outgroup. (B) Species tree confirming taxonomic identification of AF UD1. The tree was built by FastTree based on orthofinder clustering.

**Figure 3:**
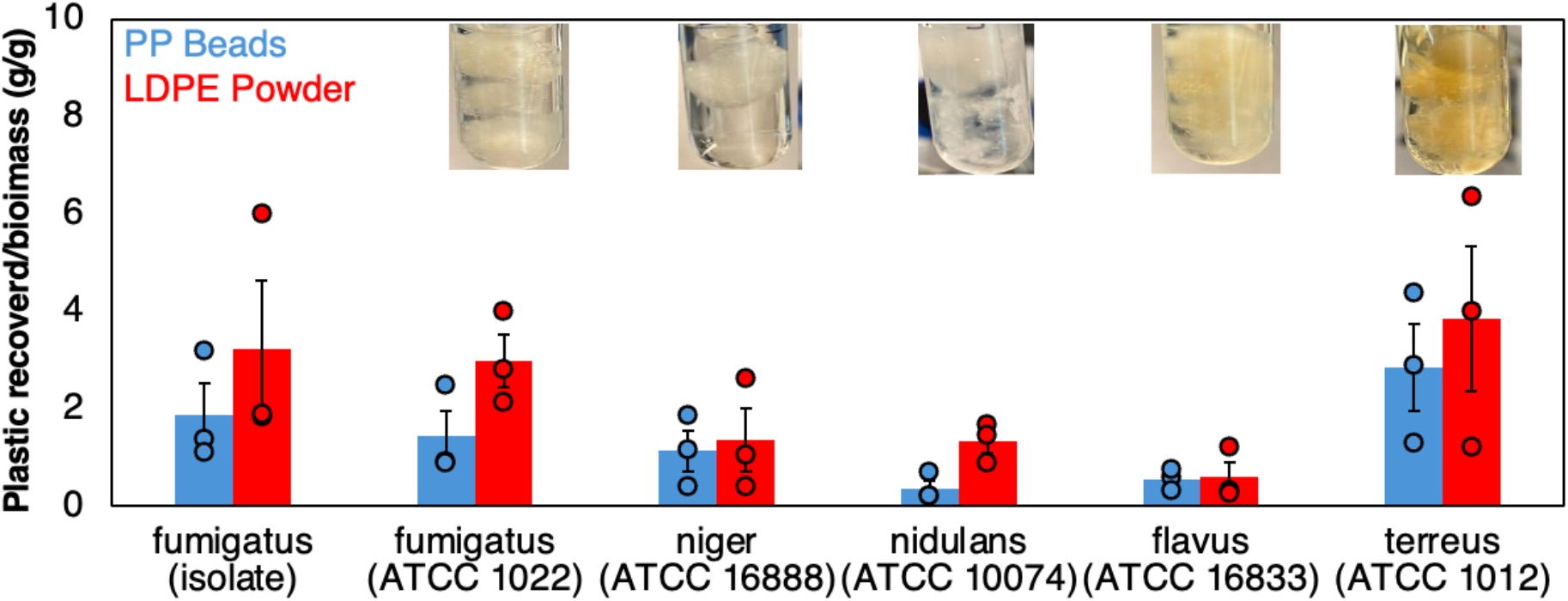
Biomass normalized flocculation of two plastic types by *Aspergillus* strains across the genome. Images above each bar are 5 mL liquid cultures of each strain with flocculated LDPE particles.

### Microplastics flocculation is driven by redox-sensitive protein interactions

Confocal and scanning electron microscope (SEM) were used to observe microplastics flocculation and better understand underlying molecular phenomena. Microplastic particles are embedded both on the AF-UD1 surface and within the hyphal network (Fig. 4A). SEM images show a dense network of hyphae and extracellular polymeric substances (EPS) that pull plastic particles into the AF-UD1 network (Fig. 4B). The dense EPS and embedded nature of the microplastics that AF-UD1 forms a stable floc that can be mechanically perturbed without a loss of plastic. The formation of robust biofilm suggests that microplastics are pulled into the fungal matrix through hydrophobic interactions with secreted or membrane bound chemicals or biomolecules produced by the fungus.

**Figure 4:**
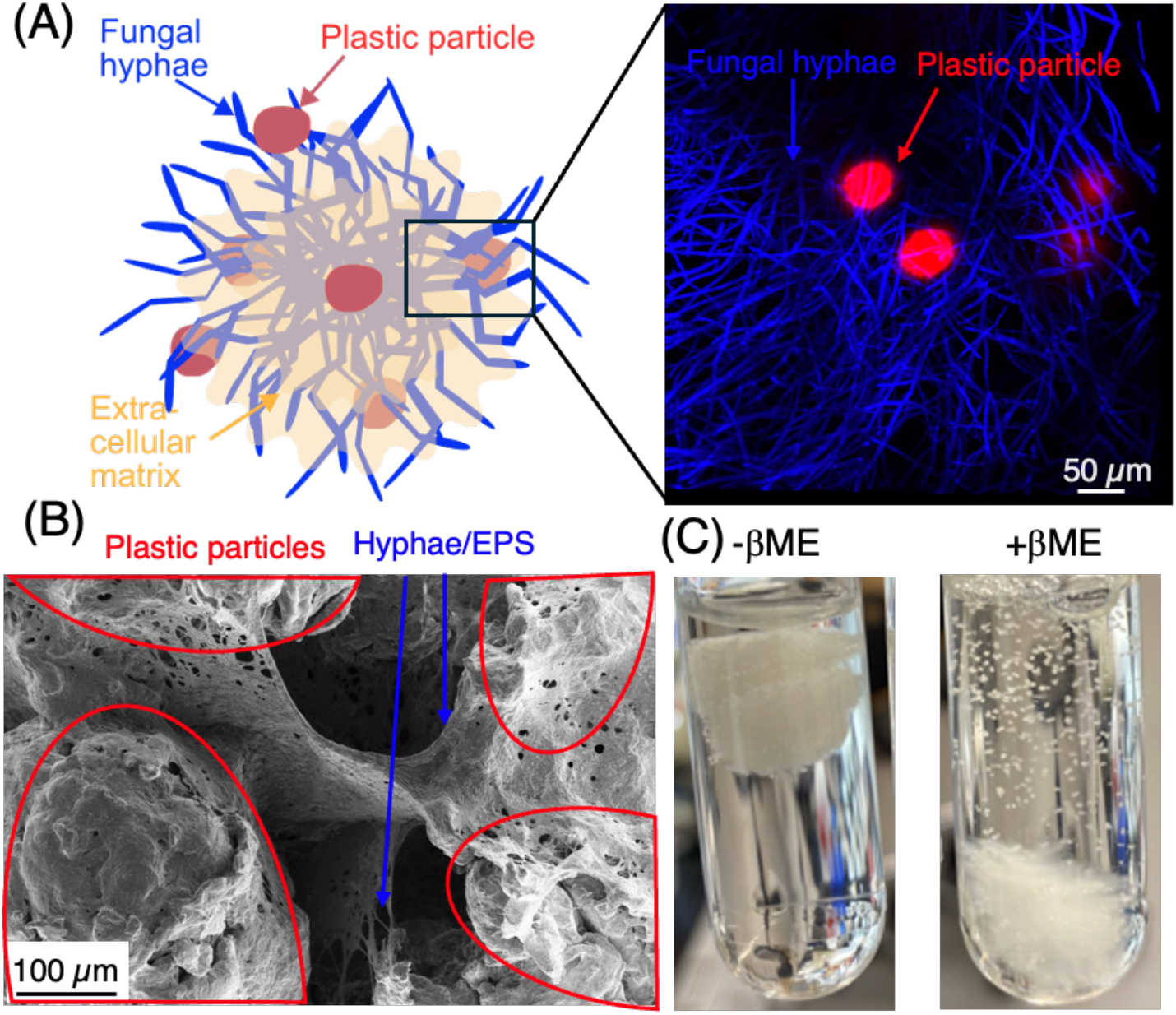
Hydrophobins drive microplastics flocculation. (A) Confocal microscopy image of *Aspergillus fumigatus* AF-UD1 (blue) stained with calcofluor white using hyphal interactions to grab fluorescent LDPE beads (red). (B) SEM image showing dense hyphal network of *Aspergillus fumigatus* AF-UD1 holding microplastic beads in a floc. (C) Images showing microplastics flocculation by AF-UD1 in the absence (left) and no flocculation in the presence (right) of beta-mercaptoethanol.

*Aspergilli* adhesion to extracellular surfaces is canonically driven by surface proteins (37). Hydrophobins are a highly surface-abundant class of proteins in *Aspergillus*, that have surfactant-like properties, namely amphiphilicity, making them very likely candidates to bind to extremely hydrophobic plastics (27, 37). Moreover, hydrophobins are predominant proteins in the outermost hydrophobic layer (27, 37) of *Aspergillus fumigatus* that form at hydrophobic/hydrophilic interfaces (38). Hydrophobins are characterized by eight conserved cysteine residues that form disulfide bonds that are responsible for stabilizing a large, hydrophobic solvent exposed interface (38). We disturbed these disulfide bonds using beta-mercaptoethanol (BME) (39), removing hydrophobins from the AF-UD1 surface, to determine if hydrophobins play a role in microplastic binding. Microplastics flocculation ability was eliminated upon the addition of BME to the culture, consistent with surface proteins such as hydrophobins that rely on disulfide bonds for structure being integral to microplastics recovery processes (Fig. 4C).

### Hydrophobins are necessary for microplastics flocculation

The role of hydrophobins in microplastics flocculation was directly assessed by repeating microplastics flocculation assays using *Aspergillus fumigatus* strains with each hydrophobin knocked out of the genome. The *Aspergillus fumigatus* genome encodes 7 different hydrophobin genes, each expressing a different hydrophobin (27). Genes for hydrophobin expression are *RODA, RODB, RODC, RODD, RODE, RODG*, and *RODF*, corresponding to proteins RodA through RodF (27). Knocking out each hydrophobin gene reduced microplastics flocculation by *Aspergillus fumigatus* (Fig. 5A). AFΔRodA, AFΔRodG, and total knockout strain AFΔRodA-G all showed statistically significant decreases in flocculation relative to wild type AF-UD1, with AFΔRodG and AFΔRodA-G failing to flocculate plastics entirely (Fig. 5A). Rod A and RodG likely play an integral role in microplastics flocculation due to the observed significant decreases. Importantly, the inhibition of microplastics flocculation by the total knockout strain (ΔRodA-G) indicates that hydrophobins are necessary for microplastics flocculation.

**Figure 5:**
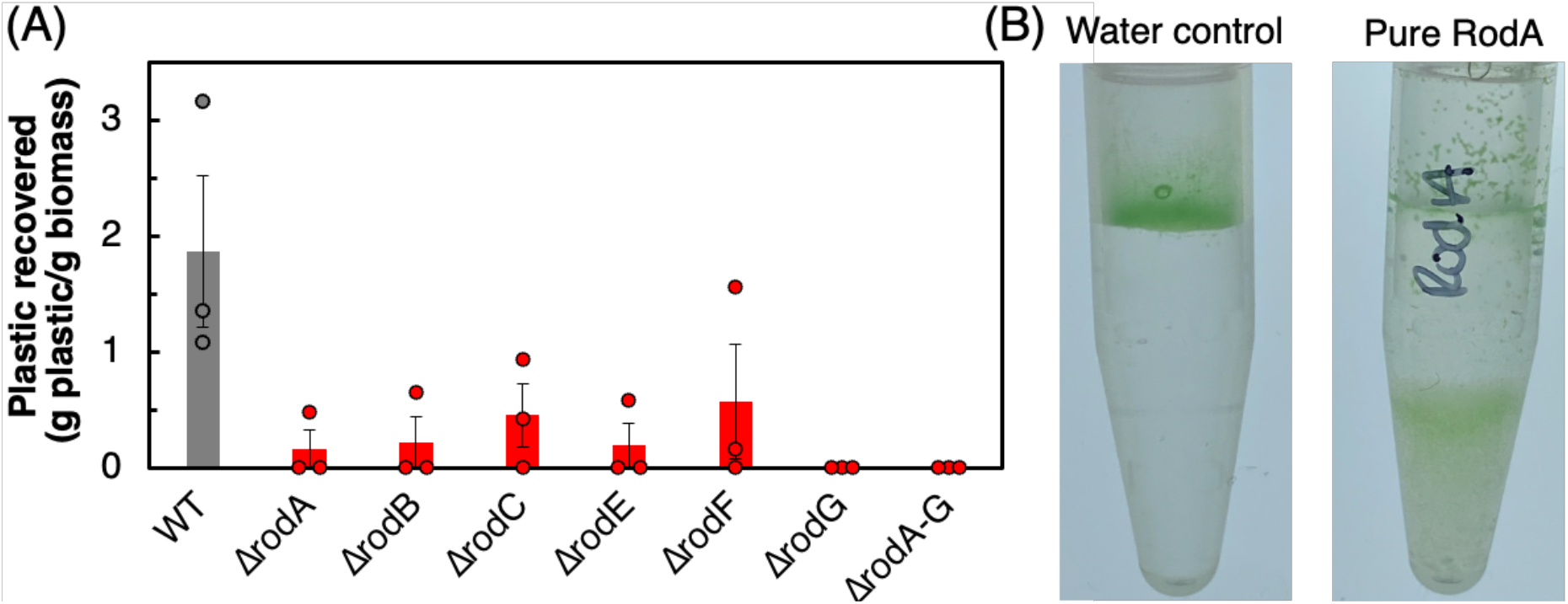
Hydrophobins are responsible for microplastics flocculation by AF-UD1. (A) Microplastics flocculation by *Aspergillus* strains with hydrophobin genes knocked out from the genome. *Aspergillus fumigatus* strains with each hydrophobin gene knocked out have reduced flocculation ability, indicating the importance of hydrophobins in microplastics recovery. (B) Recovery of green fluorescent LDPE beads by pure RodA (right) relative to a water control (left).

Hydrophobin knockout data suggest that hydrophobins are necessary for microplastics flocculation, but use of purified hydrophobin in isolation of the host is necessary to confirm their propensity for microplastic flocculation. We thus expressed RodA, reported as the hydrophobin in *A. fumigatus* responsible for cell wall surface hydrophobicity (27), in heterologous host *Yarrowia lipolytica* and subsequently purified the protein to directly assess microplastics flocculation ability by RodA in the absence of the host organism. Pure RodA flocculated microplastics from solution (Fig. 5B). Purified RodA resides in the bottom of the tube due to a density greater than that of water and microplastic particles aggregate in that area after shaking, becoming entrapped in the purified hydrophobin. Microplastics recovery by pure RodA and supporting hydrophobin knockout data demonstrate that hydrophobins are essential for microplastics flocculation by *Aspergilli*.

## Discussion

Understanding microplastics colonization is essential to mitigate potential health defects caused by pathogenic microorganisms entering the food chain through microplastic-bound biofilms. In this study, we evaluate *Aspergillus* as a model, sometimes pathogenic (27), genus of fungi to better understand fungal interactions with microplastics due to their ubiquity across soil and marine microbial communities (30, 31). We showed that *Aspergilli* are efficient microplastics binding and recovery agents. We verified that the microplastics binding phenotype is conserved across the genus by demonstrating microplastics recovery with a subset of *Aspergilli* across 5 sub-genera: *Fumigati, Nidulantes, Wentii, Terrei, and Nigri* (40). Our data suggest that microplastics binding and capture occurs independent of plastic type and size, capturing all single and mixed plastics with nearly 100% recovery, consistent with previous studies detailing 100% recovery of 200 nm PS and 5µm PMMA by *Aspergillus niger* (29). This plastic type-independent microplastics binding is consistent with reports that biofilm taxonomic composition does not vary with plastic type (41). Rather, environmental factors such as temperature, pH, and salinity drive microplastics binding interactions and dictate which taxa persist in microplastic microbial communities (18). Microplastics binding interactions require hydrophobic interactions between the microorganism and hydrophobic microplastic surface (42, 43), meaning that any mechanism altering surface hydrophobicity would be agnostic to plastic type and taxa would not change relative to polymer chemistry.

Microplastic-bound biofilms have historically been studied by identifying the dominant bacterial members present (13, 17, 19, 41), overlooking the role of the biomolecules that drive colonization and the contributions of fungal biofilm species (20, 28). We demonstrate that hydrophobins are important to microplastics binding by showing a decrease in microplastics flocculation upon knockout of each hydrophobin gene out of the genome. More importantly, we showed that pure RodA, the most abundant *A. fumigatus* hydrophobin, flocculates microplastics in isolation from the host organism, demonstrating that pure hydrophobin proteins bind directly to microplastic particles and flocculate them. The discovery of this relationship between hydrophobins and microplastics in *Aspergillus* biofilms allows for advances in pathogenicity and biodeconstruction efforts by providing the physiological context in which *Aspergilli* bind to biofilms.

Due to their inherent hydrophobicity, hydrophobins have been shown to interact with plastic substrates, namely in the context of biological deconstruction of plastics by fungal enzymes. For example, *Aspergillus oryzae* expresses hydrophobin RolA that recruits a cutinase that hydrolyses polybutylene-succinate-coadipate (44, 45). Moreover, RolA incubation with PET substrate prior to treatment with a PETase improved PET deconstruction from 17% to 26% weight loss (46). While these studies have focused on fungal enzymes and natural complexing with hydrophobins, we detail efficient microbial microplastics binding via hydrophobins. Our work highlights one strategy by which microbes colonize suspended microplastic particles, which would be the first step of biological deconstruction. Engineering this process may lead to more efficient/rapid plastics bioconstruction. For example, PET deconstruction by a PETase was improved 328-fold relative to pure PETase and 9-fold relative to surface displayed PETase by co-displaying the PETase with HFBI, a hydrophobin from *T. reesei*, on heterologous host *Pichia pastoris* (47). The work presented in this manuscript can build on such studies by providing a library of hydrophobins for plastics binding from *Aspergilli* that can be used to similarly enhance biological (micro)plastic deconstruction efforts.

Biologically compatible (micro)plastics binding technologies further enhance bioremediation efforts by providing a microplastics capture mechanism that interfaces with (bio)deconstruction efforts. Existing microplastics capture technologies used in wastewater treatment plants (WWTPs), an extremely large source of microplastics (48), such as ultrafiltration, reverse osmosis, and chemical flocculation fail to provide a mechanism through which plastics can be deconstructed. Without conversion of (micro)plastics waste into non-plastic, non-toxic products, the (micro)plastics waste crisis remains unresolved. Importantly, plastics wastes need to be upcycled into consumer products or recycled into plastics of equal value to the recycled waste to meet economic demands required to compete with plastics production from petrochemical refining (49). Hydrophobins can thus be used to capture microplastics from aqueous environments with nearly 100% efficiency and can be utilized concurrently to engineer improved biological plastics deconstruction technologies that may be able to circumvent economic barriers with conventional mechanical or chemical plastics recycling (49). Continued research on the interactions between fungal systems and microplastics is essential to identify hydrophobins capable of increased plastic binding that can ultimately be used to develop biological plastics deconstruction technologies that can mitigate the (micro)plastics waste accumulation crisis.

## Materials and methods

### Organism Isolation

*Aspergillus fumigatus* AF-UD1 was isolated from the gut of a yellow mealworm (*Tenebrio* molitor larvae) fed HDPE for 20 days. 10 mealworm guts were extracted, suspended in 1 mL of PBS, and vortexed to homogenize. Gut contents were plated on fungal Medium B (defined previously (50)). Individual colonies were re-plated on Medium B to isolate the organism. The organism was originally isolated in a co-culture with an un-identified bacterial strain. AF-UD1 was isolated from the co-culture by plating on potato dextrose agar with penicillin and streptomycin.

### Organism Identification

Whole genome sequencing was carried out on genomic DNA from AF-UD1.

### DNA Extraction

High molecular weight DNA was extracted from mycelium using the protocol of Puppo et al (2017)(51) with minor modifications. Flash-frozen biomass was ground to a fine powder in a frozen mortar with liquid nitrogen followed by very gentle extraction in 3X CTAB extraction buffer (3% CTAB (hexadecyltrimethylamonium bromide), 1.4 M NaCl, 100 mM Tris pH 8.0, 20 mM EDTA, 1% 2-mercaptoethanol) for 1h at 65 °C. The mixture was cooled down and gently extracted with 24:1 Chloroform : Isoamyl alcohol. Take out the upper phase and gently extracted with 24:1 Chloroform : Isoamyl alcohol. The aqueous phase was transferred to a new tube and 1/10th volume 3 M Sodium acetate was added, gently mixed, and DNA precipitated with iso-propanol. The sample was kept in -20 °C overnight to facilitate precipitation. DNA precipitate was collected by centrifugation, washed with 70% ethanol, air dried for 5 minutes and dissolved thoroughly in elution buffer at room temperature followed by RNAse treatment. DNA purity was measured with Nanodrop, DNA concentration measured with Qubit HS kit (Invitrogen) and DNA size was validated by Femto Pulse System (Agilent).

### Genome Sequencing

The draft genome of Aspergillus fumigatus UD1 was sequenced using PacBio Multiplexed 6-10kb Ultra-Low Input library sequenced using the REVIO. An input of 50 ng of genomic DNA was sheared to 6 kb - 10 kb using the Megaruptor® 3 (Diagenode) or g-TUBE (Covaris). The sheared DNA was treated with DNA damage repair enzyme mix, end-repair/A-tailing mix and ligated with amplification adapters using SMRTbell Express Template Prep Kit 3.0 (PacBio) and purified with SMRTbell cleanup beads. The purified ligation product was split into two reactions and enriched using 10-18 cycles of PCR using barcoded amplification oligos (IDT) and SMRTbell® gDNA Sample Amplification Kit (PacBio). Up to sixteen libraries were pooled in equimolar concentrations and the pooled libraries were size-selected using the 0.75% agarose gel cassettes with Marker S1 and High Pass protocol on the BluePippin (Sage Science). The size-selected pools were treated with DNA damage repair enzyme mix, end-repair/A-tailing mix and ligated with SMRTbell sequencing adapters, a nuclease enzyme mix and purified with SMRTbell cleanup beads. CCS data was filtered with the JGI QC pipeline to remove artifacts. CCS reads were assembled with Flye version 2.9-b1768 (https://github.com/fenderglass/Flye) and subsequently polished with two rounds of RACON version 1.4.13 (52). The mitochondrial sequence was identified based on coverage, GC content, and BLAST hits to the NCBI nt database, used to filter the CCS reads to produce non-organelle CCS, and polished with two rounds of RACON version 1.4.13 (52).

The transcriptome was sequenced using Illumina. mRNA was isolated from an input of 200 ng of total RNA with oligo dT magnetic beads and fragmented to 300 bp - 400 bp with divalent cations at a high temperature. Using TruSeq stranded mRNA kit (Illumina), the fragmented mRNA was reverse transcribed to create the first strand of cDNA with random hexamers and SuperScript™ II Reverse Transcriptase (Thermo Fisher Scientific) followed by second strand synthesis. The double stranded cDNA fragments were treated with A-tailing, ligation with NEXTFLEX UDI Barcodes (PerkinElmer) and enriched using 10 cycles of PCR. The prepared libraries were quantified using KAPA Biosystems’ next-generation sequencing library qPCR kit and run on a Roche LightCycler 480 real-time PCR instrument. Sequencing of the flowcell was performed on the Illumina NovaSeq sequencer using NovaSeq XP V1.5 reagent kits, S4 flowcell, following a 2×151 indexed run recipe. RNA-Seq reads were trimmed for artifact sequence by kmer matching (kmer=25), allowing 1 mismatch, from the 3’ end of the reads, and filtered for spike-in reads, PhiX reads and reads containing any Ns.

Quality trimming of the genome was performed using the phred trimming method set at Q6. Finally, following trimming, reads under the length threshold were removed (minimum length 25 bases or 1/3 of the original read length - whichever is longer). Filtered reads were assembled into consensus sequences using Trinity v2.12.0 (53). The genome was annotated using the JGI Annotation pipeline and made publicly available via JGI fungal genome portal MycoCosm (54).

The genome is available on JGI MycoCosm at https://mycocosm.jgi.doe.gov/AspfumUD1_1/AspfumUD1_1.info.html

### Protein Clustering and Phylogenetic Analysis

Aspergillus species below were downloaded from MycoCosm and included in orthofinder v2.55 clustering with A. fumigatus UD1 (34). Briefly, all GeneCatalog proteins were clustered into orthologous groups (orthogroups) by sequence similarity. A total of 4027 orthogroups contained a single copy from every species and were aligned to produce a species tree using default methods (32, 33). The tree file was plotted along with MycoCosm assembly and gene count metrics by phytools (55). Genome references are original publications unless sourced from AspGD (56):

A. flavus NRRL 3357 (57)

A. fischeri NRRL 181 (56)

A. fumigatus Af293 (35)

A. fumigatus A1123 (36)

A. nidulans FGSC A4 (56)

A. niger NRRL3 (58)

A. novofumigatus IBT 16806 (59)

A. terreus NIH 2624 (56)

A. udagawae IFM 46973 (60)

### Microplastics recovery assays

*Aspergillus* pre-cultures were grown in YPD at 37 °C, 220 rpm in 5 mL cultures for 2 days directly from a freezer stock. The pre-culture was then inoculated into 100 mL YPD to and grown at 37 °C, 100 rpm in a 500 mL flask to grow *aspergillus* ‘flocs’ LDPE (LDPE particles were purchased from Goodfellow Cambridge Limited, Huntingdon, England; catalog number LS563303), PP (PP was obtained from post-consumer yogurt Chobani yogurt cups. For use in this study, PP disks/beads cut out using a 2 mm diameter hole punch), PET (PET was obtained from post-consumer Dasani water bottles. For use in this study, PET disks/beads cut out using a 5 mm diameter hole punch), or UHMWPE (Ultra-high molecular weight PE (UHMWPE) was purchased from Sigma-Aldirch Chemical Company; catalog number 43272 – 100g) Microplastics particles at approximately 25 mg were suspended in 5 mL of sterile mineral media (1g NaH_2_PO_4_, 0.5 g MgSO_4_*7H_2_O, 0.2 g KH_2_PO_4_, and 1 g yeast extract per 1 liter). 2-3 flocs from the large *aspergillus* culture were dropped into the 5 mL culture containing plastic powder. The cultures were allowed to shake at 30 °C, 220 rpm to allow the aspergillus strains to slowly grow in a nutrient deprived environment. Every 2 hours, the culture was shaken to allow fungi to come into direct contact with the plastic. Once plastic was grabbed at one of these 2 hour intervals, the culture was removed for analysis. If no plastic was grabbed by the fungi (in the case of aspergillus knockout experiments) after 36 hours, the culture was removed and discarded.

For mass-normalized microplastics recovery assays, flocs of *A. fumigatus* UD01, *A. fumigatus* (ATCC 1022) *A. niger* (ATCC 16888), *A. nidulans* (ATCC10074), *A. flavus* (ATCC 16833), or *A. terreus* (ATCC 1012) containing microplastics were removed from the culture tubes and allowed to dry. Remaining plastic was subsequently removed from the tube, dried, and weighed. Mass of flocculated plastic was calculated by subtracting the mass of remaining plastic from the initial plastic mass. Additionally, the microplastic-flocculated cultures were weighed. Calculated flocculated plastic mass was subtracted from the mass of the dried flocculated plastic fungal culture to obtain dry biomass weight.

For microplastics recovery assays that included beta-mercaptoethanol (BME), the BME was added into the culture tube with plastic and prior to adding the fungal flocs. Flocs were added and the protocol outlined above was followed.

For recovery assay with pure RodA, 10 mg of green fluorescent LDPE particles (Green LDPE beads were purchased from Cospheric LLC, Somis, CA; catalog number UVPMS-BG-1.00 35-45 um, respectively) were placed into 1 mL of solution containing purified RodA. The solution was lightly shaken to mix and then left stationary at room temperature overnight to allow separation.

### Confocal microscopy

Microplastics recovery assays were carried out as above, but with red fluorescent LDPE beads (Red Fluorescent LDPE beads were purchased from Cospheric LLC, Somis, CA; catalog numbers UVPMS-BR-0.995 45-53 um). After flocculation, floc culture was washed with PBS three times. The culture was placed into a microscopy sample dish (ibidi ibiTreat: #1.5 polymer coverslip, tissue culture treated, sterilized) in 2 mL PBS. One microliter of calcofluor white was added to the culture and it was stored in darkness for 15 minutes to stain the fungi. Microscopy images were taken using the NIIMBL Stellaris 8 tauSTED/FLIM Confocal Microscope at the University of Delaware Bioimaging Center.

### Scanning electron microscopy

Microplastics recovery assays were carried out as above. After flocculation, floc culture was washed with PBS three times. Culture flocs were coated in platinum and imaged on the Apreo VolumeScope™ Scanning Electron Microscope at the University of Delaware Bioimaging Center.

### Hydrophobin knockout strains

Mutant hydrophobin knockout strains were generously donated by Jean-Paul Latgé and Isabelle Mouyna from Aspergillus Unit, Institut Pasteur, 75015 Paris, France. The hydrophobin knockouts were generated from methods listed in the original publication(27).

### RodA cloning in *Yarrowia lipolytica*

RodA sequence was codon optimized for *Yarrowia lipolytica* and the resulting gene fragment was ordered from Twist Biosciences with an AscI cutsite on the N terminus and an NheI cutsite on the C terminus. Gene fragment was digested with AscI and NheI enzymes at 37 °C for 1 hour; restriction enzymes were heat inactivated at 80 °C for 20min. The vector for cloning was a homology donor for integration into the AXP site in Yarrowia lipolytica(61). The vector was also digested with AscI and NheI at 37 °C for 1 hour. The vector and insert were ligated using NEB DNA ligase. 5uL of ligation mix was added to 50uL of NEB 10β competent cells which were heat shocked at 42 °C for 30 seconds and then recovered in 1mL of LB media for 1 hour at 37ºC with shaking. 100uL of transformed cells were plated onto an ampicillin containing LB plate and grown overnight at 37 °C. The sequence of the resulting plasmid was confirmed by Sanger sequencing. The RodA gene was integrated into the AXP site using the homology donor and a CRISPR containing plasmid. Integration was confirmed via colony PCR and then the strain was cured of all plasmids.

### Codon optimized RodA sequence

ATGAAGTTTAGCCTCTCTGCTGCAGTACTGGCCTTTGCCGTGTCTGTGGCTGCGCTCCCCCAGCACGATGTCAACGCCGCTGGAAACGGTGTCGGCAACAAAGGCAATGCCAACGTGCGAT TCCCTGTTCCCGACGACATCACCGTTAAACAAGCAACTGAGAAGTGTGGAGACCAGGCCCAGCTGTCATGCTGCAACAAGGCCACCTACGCTGGCGACGTGACGGATATCGACGAGGGTATTCTGGCGGGTACTCTCAAGAACCTCATCGGCGGGGGCTCGGGAACAGAAGGACTAGGTTTGTTCAACCAGTGTTCCAAGCTGGATCTGCAGATTCCTGTCATTGGCATCCCCATCCAGGCTCTTGTTAACCAAAAGTGCAAGCAGAACATAGCCTGTTGCCAGAATTCGCCGTCCGACGCCAGTGGCTCTCTGATTGGACTTGGTCTTCCATGTATTGCTCTGGGATCCATCTTGTAG

### RodA protein sequence

MKFSLSAAVLAFAVSVAALPQHDVNAAGNGVGNKGNANVRFPVPDDITVKQATEKCGDQAQLSCCNKATYAGDVTDIDEGILAGTLKNLIGGGSGTEGLGLFNQCSKLDLQIPVIGIPIQALVNQKCKQNIACCQNSPSDASGSLIGLGLPCIALGSIL

### RodA expression and purification

*Yarrowia lipolytica* RodA was grown for 4 days at 28ºC with agitation at 220 rpm in 50 mL YPD. Culture was transferred to 50mL falcon tube and centrifuged at 4000 rpm to separate pellet and supernatant. Supernatant was ultra-centrifuged for 1hr at 100,000g in SW32Ti rotor using OptiSeal 32mL tubes and adapters from Beckman Coulter. Supernatant was removed and pellet was resuspended in 1 mL of 2% SDS. Sample was transferred to microcentrifuge tube and boiled for 10 min at 98 °C. In Beckman Coulter’s 5mL thinwall open top tubes, sample was ultra-centrifuged at 100,000 g in 5 mL of SDS for 1 hr at 20 °C using SW55Ti rotor. This process was repeated 2 times. All SDS was removed and the pellet was resuspended in 5mL DI water. The sample was then ultra-centrifuged at 100,000 g for 1 hr at 20 °C using SW55Ti rotor. This process was repeated 2 times. Resulting hydrophobin pellet was transferred to a 1.5 mL microcentrifuge tube with 1mL DI water. RodA expression and purification protocol was adapted from(62).

## Acknowledgements

This research was funded in part by the Chemistry Biology Interface at the University of Delaware, under NIH training grant T32GM133395. This research was funded in part by the Delaware Environmental Institute, University of Delaware. Reagents for this research were ordered in part with funds from the QIAGEN Young Scientist Research Grant award 2022. This material is based upon work supported by the U.S. Department of Energy, Office of Science, Office of Biological and Environmental Research under Award Numbers DE-SC0022018 and DE-SC0023085. The work (proposal: 10.46936/fics.proj.2021.60038/60000396) conducted by the U.S. Department of Energy Joint Genome Institute (https://ror.org/04xm1d337), a DOE Office of Science User Facility, is supported by the Office of Science of the U.S. Department of Energy operated under Contract No. DE-AC02-05CH11231.

A portion of this research was performed under the Facilities Integrating Collaborations for User Science (FICUS) program (proposal: 508042) and used resources at the DOE Joint Genome Institute (https://ror.org/04xm1d337) and the Environmental Molecular Sciences Laboratory (https://ror.org/04rc0xn13), which are DOE Office of Science User Facilities operated under Contract Nos. DE-AC02-05CH11231 (JGI) and DE-AC05-76RL01830 (EMSL). Sequencing

Project ID: 1428900. Final Deliverable Project ID: 1428895. Microscopy access was supported by grants from the NIH-NIGMS (P20 GM103446), the NIGMS (P20 GM139760) and the State of Delaware. Leica Stellaris 8 tauSTED/FLIM Confocal was acquired with NIST (70NANB21H085). We thank Debbie Powell and Sylvain Le Marchand for exceptional support on scanning electron microscopy and confocal microscopy, respectively. We also acknowledge Jenna Moore-Ott for the generation of cartoons that led to creation of figures in this work.

## Author Contributions (CRediT)

Ross R. Klauer: led all experimental work, investigation, methodology, validation, formal analysis, writing – original draft

Rachel Silvestri: investigation, methodology, writing – review and editing

Hanna White: investigation, writing – review and editing

Richard Hayes: methodology

Robert Riley: methodology

Anna Lipzen: methodology

Kerrie Barry: coordination

Igor V. Grigoriev: coordination

Jayson Talag: methodology

Victoria Bunting: methodology

Zachary Stevenson: investigation

Kevin Solomon: conceptualized the study, supervised the study, acquired funding, and reviewed & edited the manuscript.

Mark Blenner: conceptualized the study, supervised the study, acquired funding, and reviewed & edited the manuscript.

